# Neural Correlates of the Embodied Sense of Agency

**DOI:** 10.1101/2024.09.06.611665

**Authors:** Amit Regev Krugwasser, Reina van der Goot, Geffen Markusfeld, Yair Zvilichovsky, Roy Salomon

## Abstract

The Sense of Agency (SoA) is the subjective experience that ‘I am in control of my actions’. Recent accounts have distinguished two levels in the formation of SoA - early implicit sensorimotor processes (feeling of agency) and later explicit higher-level processes, incorporating one’s thoughts and beliefs (judgment of agency). Even though SoA is fundamental to our interactions with the external world and the construct of the self, its underlying neural mechanism remains elusive. In the current pre-registered electroencephalography (EEG) study, we used time-frequency analysis and Multivariate Pattern Analysis (MVPA) to investigate the electrophysiological characteristics associated with SoA. We used an established embodied virtual reality paradigm in which visual feedback of a movement is modulated to examine the effect of conflicts between the predicted and perceived sensory feedback. Participants moved their finger and were shown a virtual hand that either mimicked their movement or differed anatomically (different finger) or spatially (angular shift), then judged their SoA over the observed movement while their brain activity was recorded. In accordance with our pre-registered hypothesis, we found that a reduction of SoA is associated with decreased attenuation in the alpha frequency band. Increased power in the theta frequency band was also associated with SoA reduction. Importantly, we show that trials containing a sensorimotor alteration vs. trials containing no alteration can reliably be decoded with up to 68% accuracy starting around 200ms after the movement onset. Finally, cross-decoding analyses revealed similar neural patterns for reduced SoA in the anatomical and spatial conditions, starting around 500ms after the movement onset. Together, our results reveal a cortical signature of loss of SoA and provide neural evidence supporting the hypothesis of a two-level formation of SoA - an early domain-specific component, possibly the equivalent of the implicit feeling of agency, and a late domain-general component, possibly the equivalent of the explicit judgment of agency.

## Introduction

The Sense of Agency (SoA) is the feeling of control over one’s actions (Haggard et al., 2002). It allows us to sense ourselves as active embodied agents who can influence the world around us and shape our lives (Gallagher, 2000), enabling the distinction between ourselves and the environment (Knoblich et al., 2003; Krugwasser et al., 2022; Synofzik et al., 2010). Despite its centrality in shaping one’s bodily-self, it is considered phenomenologically “transparent”, rarely reaching conscious awareness in day-to-day scenarios. However, violations of SoA are quite salient and often experienced consciously (Krugwasser et al., 2019; Stern et al., 2022; Tapal et al., 2017). For example, if I were to make a movement with my right hand and my left hand moved instead, this would cause a disruption of SoA, a phenomenon found in numerous neurological and psychiatric conditions (Assal et al., 2007; Harduf et al., 2023; Krugwasser et al., 2022; Moore & Fletcher, 2012; Moro et al., 2015). While SoA can extend to action outcomes in general, e.g., the experience of instrumental control over an external object or event (Gozli & Brown, 2011; Haggard, 2017; Nakashima, 2019), we suggest that the core mechanism is based upon embodied SoA, the attribution of authorship of action to oneself (Pyasik et al., 2019; Wen et al., 2023).

Several mechanisms have been suggested to explain the formation of SoA. For example, the “Comparator Model” suggests that actions are continuously assessed by comparing the predicted sensory outcomes (forward models) and the actual (Applebaum et al., 2024; Engbert et al., 2008; Haggard, 2017; Krugwasser et al., 2022; Wolpert et al., 1995; Zaidel & Salomon, 2023). Computationally, the forward model comprises specific predictions related to various action characteristics, such as spatial, anatomical, and temporal aspects (Synofzik et al., 2013; Tsakiris et al., 2005; Villa et al., 2018). For instance, a reaching movement generates predictions regarding the timing of visual feedback (seeing the hand move exactly when it moves), the anatomical limb involved (which hand moves), and the direction of the movement, as well as prediction regarding tactile, proprioception, and auditory sensations. We have previously shown that the sensitivity to conflicts in spatial, anatomical, and temporal aspects of action is highly correlated (Krugwasser et al., 2019). Participants who performed well in one sensorimotor conflict – usually performed well in other conflicts as well. Decision criterion was also positively correlated between these aspects, suggesting a central decisional mechanism of SoA for predictions across different aspects of movement.

Another theory of SoA is the theory of apparent mental causation (Wegner & Wheatley, 1999). This theory posits that SoA is based on three factors: Priority, consistency, and exclusivity. Priority relates to the proximity in time between one’s intention to act and the action itself, with the intention preceding the action. Consistency refers to the degree of correspondence between the content of one’s intention and the actual action performed. Exclusivity pertains to the absence of other perceived causes for the action. The theory maintains that if any of these factors is absent, SoA is reduced (Braun et al., 2018; Moore, 2016; Villa et al., 2022).

Despite their fundamentally different natures, with the Comparator Model being predictive and the theory of apparent mental causation being post-deductive, it is possible they are complementary. The two-step theory (Synofzik et al., 2008) combined these by suggesting SoA is based on two distinct processes: (1) A perceptual level, characterized by a non-conceptual, low-level Feeling of Agency. This feeling distinguishes between self-caused and non-self-caused actions, without assigning the cause to a specific external entity. Similar to the Comparator Model, the Feeling of Agency is thought to be based on sensorimotor integration, where internal predictions and sensory feedback are weighed and compared. (2) A conceptual Judgment of Agency, responsible for making a higher-level attribution of agency. The Judgment of Agency mechanism processes the pre- conceptual Feeling of Agency within a framework that incorporates one’s beliefs and prior information, to attribute agency to either oneself or to a specific external entity. When the internal prediction and sensory feedback are congruent, a feeling of a self-generated movement is established (Feeling of Agency). However, when a conflict occurs, the sensorimotor prediction error propagates to a higher-level mechanism (Judgment of Agency) which attempts to explain the discrepancy. This approach bridges the gap between predictive and post-deductive theories, resolving the apparent conflict between them: When an action’s outcome aligns closely with the prediction, generating a minimal prediction error (or none), low-level, implicit sensorimotor processing is sufficient. In contrast, when the action outcome diverges from the predicted outcome, generating a more substantial prediction error, low-level processing might not be enough and additional high-level, explicit, processing will be required to resolve the conflict (Leptourgos & Corlett, 2020). For example, when we reach for a cup, despite the presence of sensory and neural noise, our low-level sensorimotor systems typically suffice to do this effortlessly. However, if snag our sleeve hindering our movement, the reaching action may not align closely with our predictions. This discrepancy generates a substantial prediction error, as the actual outcome deviates from what was predicted. In such a situation, low-level processing alone might not be enough to correct the action. Additional high-level processing is required to quickly assess the situation and adjust the action (e.g., unsnag our sleeve by moving hand backwards). The two-step theory bridges the predictive and post-deductive perspectives into a conceptually coherent framework, however empirical evidence is scarce and the neural correlates of these mechanisms are not well understood.

Most studies relating to the neural mechanisms of SoA have relied on non-embodied paradigms in which the contingencies between actions and outcomes are learned during the experiment (e.g. Fukushima et al., 2013; Ohata et al., 2020; Renes et al., 2015; Tsakiris et al., 2010). SoA processes were shown to be connected to activation in the temporoparietal junction (TPJ, see Balslev et al., 2006; Farrer et al., 2008; Fukushima et al., 2013; Spengler et al., 2009), a region that is known to integrate multisensory body-related information (Arzy et al., 2006; Blanke, 2012; Quesque & Brass, 2019). Other brain regions related to SoA include the pre-SMA (Supplementary Motor Area, see Kühn et al., 2013; Miele et al., 2011; Yomogida et al., 2010), which has a key role in preparing and initiating voluntary actions (Seghezzi & Zapparoli, 2020), and the Cerebellum (Agnew & Wise, 2008; de Bézenac et al., 2016; Leube et al., 2003), which monitors self-motion information (Andre et al., 2024; Rondi-Reig et al., 2014). These motor regions were suggested to be a part of a neural circuitry representing the “Comparator Model” (Haggard, 2017; Harduf et al., 2023). The insula, involved in interoceptive processing associated with self-awareness and higher-order self-referential processing (Frewen et al., 2020; Park et al., 2018), was also found to take part in SoA formation (Harduf et al., 2023; Karnath & Baier, 2010; Leube et al., 2003). In addition, in an intracranial study that used an embodied paradigm, the primary motor cortex (M1) was shown to take part in processing sensorimotor conflicts (Serino et al., 2022), a central mechanism in SoA formation (Krugwasser et al., 2019; Ohata et al., 2020; Synofzik et al., 2008). Collectively, these findings suggest a distributed neural network underlying SoA formation, highlighting the complexity of the cognitive processes involved in attributing agency to one’s actions. While these findings are relatively consistent (see Seghezzi et al., 2019; Zito et al., 2020 for meta-analyses) and provide important information regarding the brain regions involved in SoA formation, little is known regarding the timing of embodied SoA processing. Despite its importance, the electrophysiological attributes of the neural mechanisms underlying SoA are still not well understood. Only a handful of studies touched upon the electrophysiological mechanisms of SoA, and even fewer have examined embodied SoA, in which our priors are stronger due to years of experience with our body.

Event-related potential (ERP) studies of SoA revealed that self-generated actions are followed by a smaller ERP amplitude compared to non-self-generated actions, for both visual (Desantis et al., 2014; Hughes & Waszak, 2011) and auditory effects (Horváth, 2015; Mifsud et al., 2016), as early as 100ms following stimuli onset. A more recent study showed the same effect early after movement onset (70-135ms) but the opposite, namely a higher amplitude in self-generated actions, at a later time (165-200ms, see Han et al., 2021). In the frequency domain, activity in the alpha band (8-12 Hz) was found to be attenuated when there was a close match between the intended and observed action, using a task in which the visual consequences of the intended action were manipulated (Pezzetta et al., 2018). Further studies showed a strong relation between SoA and alpha band in the same direction, i.e., reduced SoA is associated with increased alpha activity (Bu-Omer et al., 2021; Kang et al., 2015; Savoie et al., 2018; Zama et al., 2019). Another frequency band that was shown to be related to SoA is low beta (13-20 Hz), in which higher power occurred when the visual feedback of a hand movement was delayed (Bu-Omer et al., 2021) or when a presented cursor trajectory deviated from the actual hand trajectory (Savoie et al., 2018). Results from the same study also showed a larger power increase in low theta (2-4 Hz) for the same comparison. This finding is in line with previous studies examining deviations from expected action effects (Mas-Herrero & Marco-Pallarés, 2014; Pezzetta et al., 2018; Tzur & Berger, 2007; although one should keep in mind that theta is defined differently by different researchers, ranging from 2-8 Hz, see Newson & Thiagarajan, 2019 for a review).

While many of the imaging studies of SoA focused on prediction violation in a single aspect of an action (e.g., unexpected tone, delayed visual feedback), real-life actions are comprised by more than one aspect of motion (i.e. temporal, spatial and anatomical). Computationally it is likely that distinct predictions are formed during action for each aspect. For example, the forward models predicting the sensory outcomes for direction of a movement and the limb in which this movement is enacted require different types of information. By elucidating how discrepancies in different aspects affect agency perception, we can potentially identify mechanisms pertaining to specific prediction versus more supramodal representations of SoA (Harduf et al., 2023; Krugwasser et al., 2019). To do so, we need to explore the interplay between low and high-level mechanisms of SoA – Are there distinct low-level mechanisms for each sensorimotor aspect that function similarly, or is it the high-level mechanism that dictates our SoA over an action based on a unified low-level mechanism?

In the current pre-registered EEG study, we used time-frequency analysis and Multivariate Pattern Analysis (MVPA) to investigate the electrophysiological characteristics associated with embodied SoA. Using an established embodied SoA virtual reality paradigm, we manipulated the visual feedback of finger movements to examine the effect of congruency between expected and actual sensory feedback (Krugwasser et al., 2019, 2022; Stern et al., 2022). Participants were presented with either an unaltered movement, an anatomical alteration (different finger), or a spatial alteration (angular deviation) and were asked to indicate their SoA over the observed movement, while their brain activity was recorded using EEG. Our pre-registered hypotheses were: 1) SoA will decrease when anatomical or spatial alterations are introduced, and both sensitivity and bias will be correlated between these conditions. 2) Embodied SoA will be associated with alpha attenuation, i.e., a smaller power decrease in the alpha frequency band in the altered conditions compared to the unaltered condition. 3) A modulation of high beta band power when SoA is reduced. 4) Above chance-level decoding for unaltered vs. anatomical and unaltered vs. spatial conditions. Pre-registration is available at osf.io/27a9c

## Results

### Behavioral Analysis

SoA ratings drastically decreased between the Unaltered condition (0.94 ± 0.06) and either one of the altered conditions (Anatomical: 0.02 ± 0.02, Spatial: 0.1 ± 0.1), in which visual feedback was manipulated (Anatomical: t(29) = 71.12, p < 0.0001, d = 14.36; Spatial: t(29) = 32.17, p < 0.0001, d = 5.87; see Figure 1C), indicating that our experimental design successfully manipulated the subjective experience of agency. Confidence judgments showed a significant difference between the Unaltered (2.41 ± 0.5) and the Anatomical (2.76 ± 0.4) conditions (t(29) = 3.99, p < 0.001) but no significant difference between the Unaltered and the Spatial (2.36 ± 0.55) conditions (t(29) = 0.72, p = 0.47, see Supplementary Table S1). Average finger movement time (Unaltered: 742 ± 141, Anatomical: 718 ± 166, Spatial: 740 ± 170, all in ms) did not significantly differ between conditions (all p-values > 0.58). In line with our pre-registered hypothesis, both sensitivity (d’Anatomical = 4.04 ± 0.6, d’Spatial = 3.35 ± 1.05) and bias (cAnatomical = -0.27 ± 0.32, cSpatial = 0.07 ± 0.31) were highly correlated between the Anatomical and the Spatial conditions (rSensitivity = 0.84, p < 0.001; rBias = 0.5, p < 0.01).

**Figure 1.**
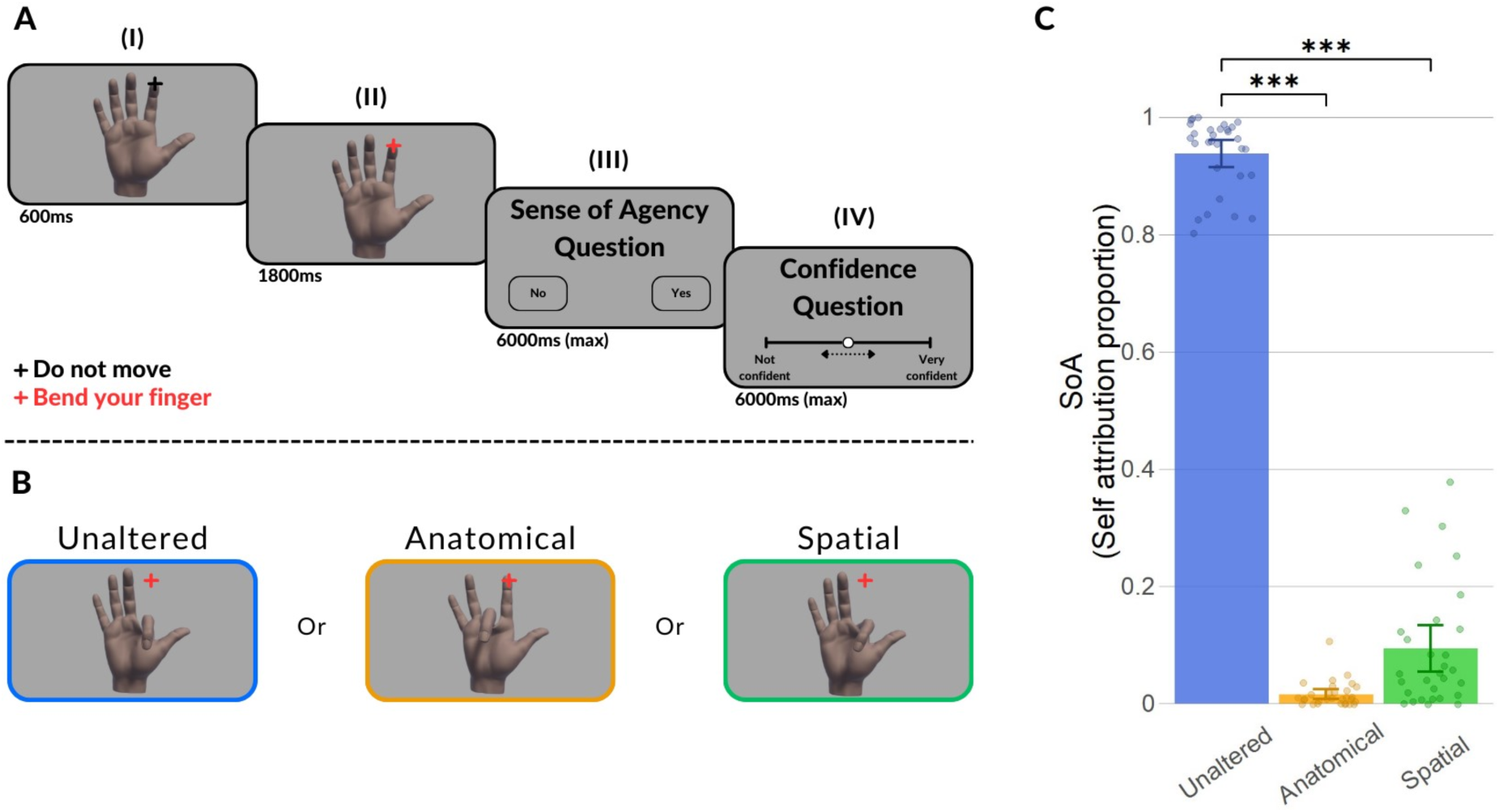
Trial flow, conditions, and behavioral results. **A**: Each trial began with the virtual hand presented, overlaid with a black fixation cross (signaling not to move yet) (I). When the fixation cross changed color to red (action cue), participants conducted a single bending movement with their index finger (II). Next, participants indicated if they had SoA over the observed movement (III) and rated their confidence regarding the SoA question (IV). **B**: Conditions (virtual hand manipulations) used in the experiment. **C**: Average self-attribution in each condition. Dots represent the average rating per individual.

### Time-Frequency Analysis

Upon establishing that subjective SoA ratings decreased dramatically for both the Anatomical and Spatial conflict conditions, we proceeded to investigate the electrophysiological power modulations associated with SoA disruptions. To this end, we followed the pre-registration and, subsequently, averaged power in both the alpha (8-12 Hz) and high beta (20-30 Hz) frequency band in our predefined temporal and spatial regions of interest (ROI) of 100ms to 500ms, for each participant, for electrodes C3, C4, and Cz.

In line with our pre-registered hypothesis, we observed a decrease in alpha attenuation in the Anatomical condition compared to the Unaltered condition in all three pre-registered electrodes, C3: t(29) = 2.51, p < 0.01, d = 0.47; C4: t(29) = 2.01, p < 0.05, d = 0.37; Cz: t(29) = 3.41, p < 0.001, d = 0.63 (see Figure 2). Contrary to the robust difference across predefined electrodes in alpha frequency band, a significant difference in high beta frequency band was only found in electrode Cz: t(29) = 2.11, p < 0.05, d = 0.39, but not in electrodes C3 (t(29) = -1.62, p = 0.11, d = 0.3, BF10 = 0.63) and C4 (t(29) = -1.6, p = 0.12, d = 0.29, BF10 = 0.6); see Supplementary Figure S1). Bayesian analysis showed inconclusive evidence for a null effect in these electrodes.

**Figure 2.**
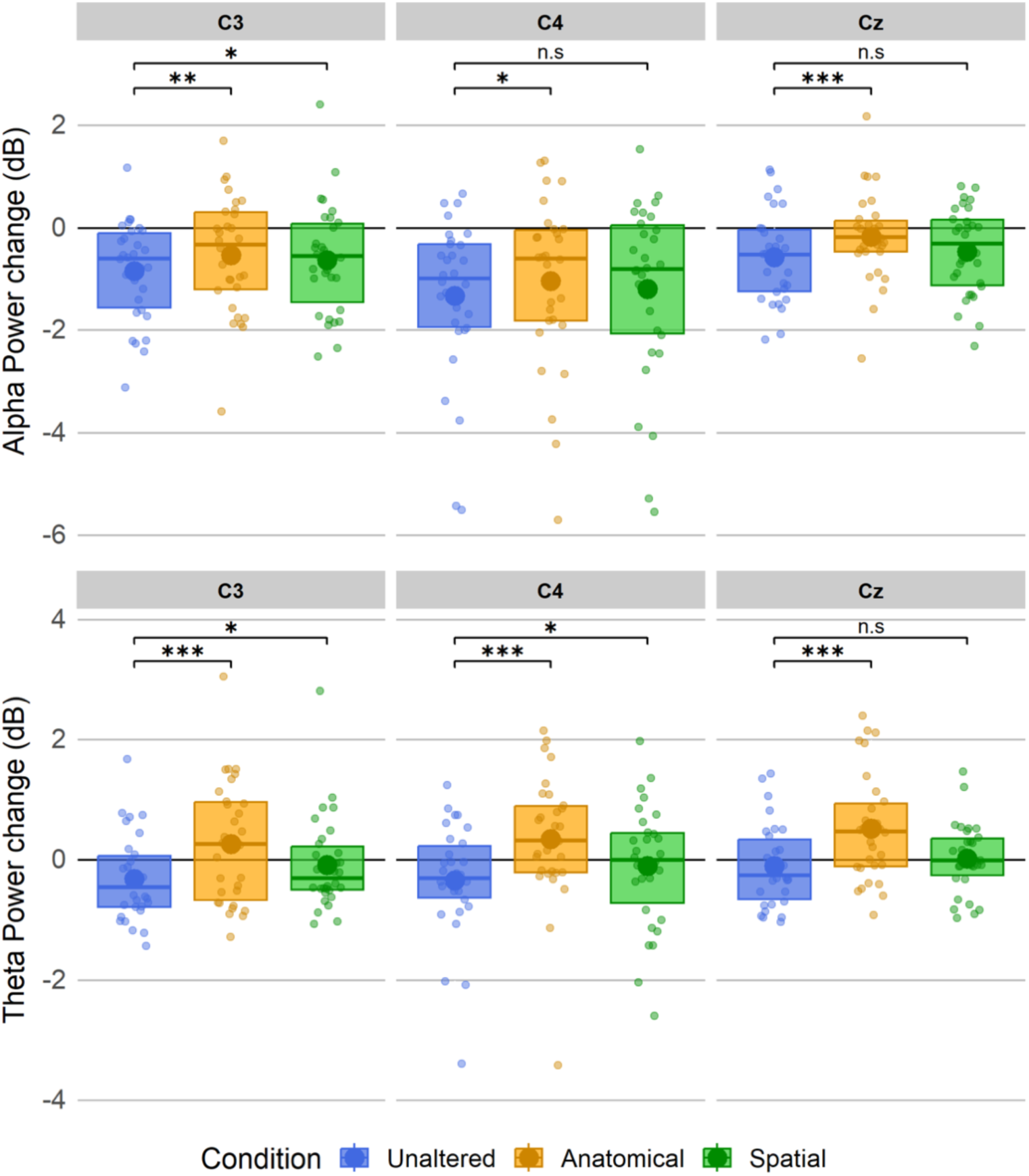
Differences in Power for pre-registered electrodes. Average (large circles) and individual participants’ (dots) power modulation for each electrode (C3, C4 & Cz) in each condition (Unaltered, Anatomical & Spatial), in alpha (top) and theta (bottom) frequency bands, compared to baseline.

Contrary to our expectations, a significant difference between the Unaltered and Spatial conditions in alpha frequency band in the pre-registered electrodes was only observed in C3: t(29) = 2.13, p < 0.05, d = 0.4 (see Figure 2). No significant difference was found in either C4 (t(29) = 1.16, p = 0.13, d = 0.21, BF10 = 0.36) or Cz (t(29) = 1.37, p = 0.09, d = 0.25, BF10 = 0.45). As before, Bayesian analysis showed inconclusive evidence for a null effect in these electrodes. No significant difference in power was found for higher beta frequency band (C3: t(29) = 0.98, p = 0.38, d = 0.18, BF10 = 0.3; C4: t(29) = 0.04, p = 0.97, d = 0.007, BF10 = 0.19; Cz: t(29) = 0.62, p = 0.54, d = 0.11, BF10 = 0.23; see Supplementary Figure S1). The subsequent Bayesian analysis showed evidence supporting the null effect.

Having identified a consistent relationship between power in the alpha frequency band and SoA disruptions within our temporal and spatial ROI across pre-registered electrodes, we examined the power distribution across time by performing non-parametric permutation statistics across the entire frequency range. For the Anatomical condition, frequency maps demonstrated that the significant clusters aligned with our hypothesized temporal ROI of 100ms to 500ms. Interestingly, all three pre-registered electrodes demonstrated that the observed decrease in alpha attenuation was part of a larger cluster ranging from theta to alpha (see Figure 3A). Electrode C3 demonstrated a cluster ranging from 180ms to 640ms from 4 to 14 Hz (p < 0.001), electrode C4 demonstrated two significant clusters, an early one ranging from 95ms to 640ms from 4 to 14 Hz (p < 0.01) and a later one ranging from 417ms to 640ms from 17 to 40 Hz (p < 0.01). Lastly, electrode Cz yielded one cluster ranging from 140ms to 640ms and 4 to 17 Hz (p < 0.001). All clusters demonstrated reduced alpha and theta attenuation, except the late high beta cluster, which demonstrated the opposite effect, with greater high beta attenuation associated with the Anatomical condition. In contrast to the Anatomical condition, no significant clusters were found for the Spatial condition using non-parametric permutation analyses (see Figure 3B).

**Figure 3.**
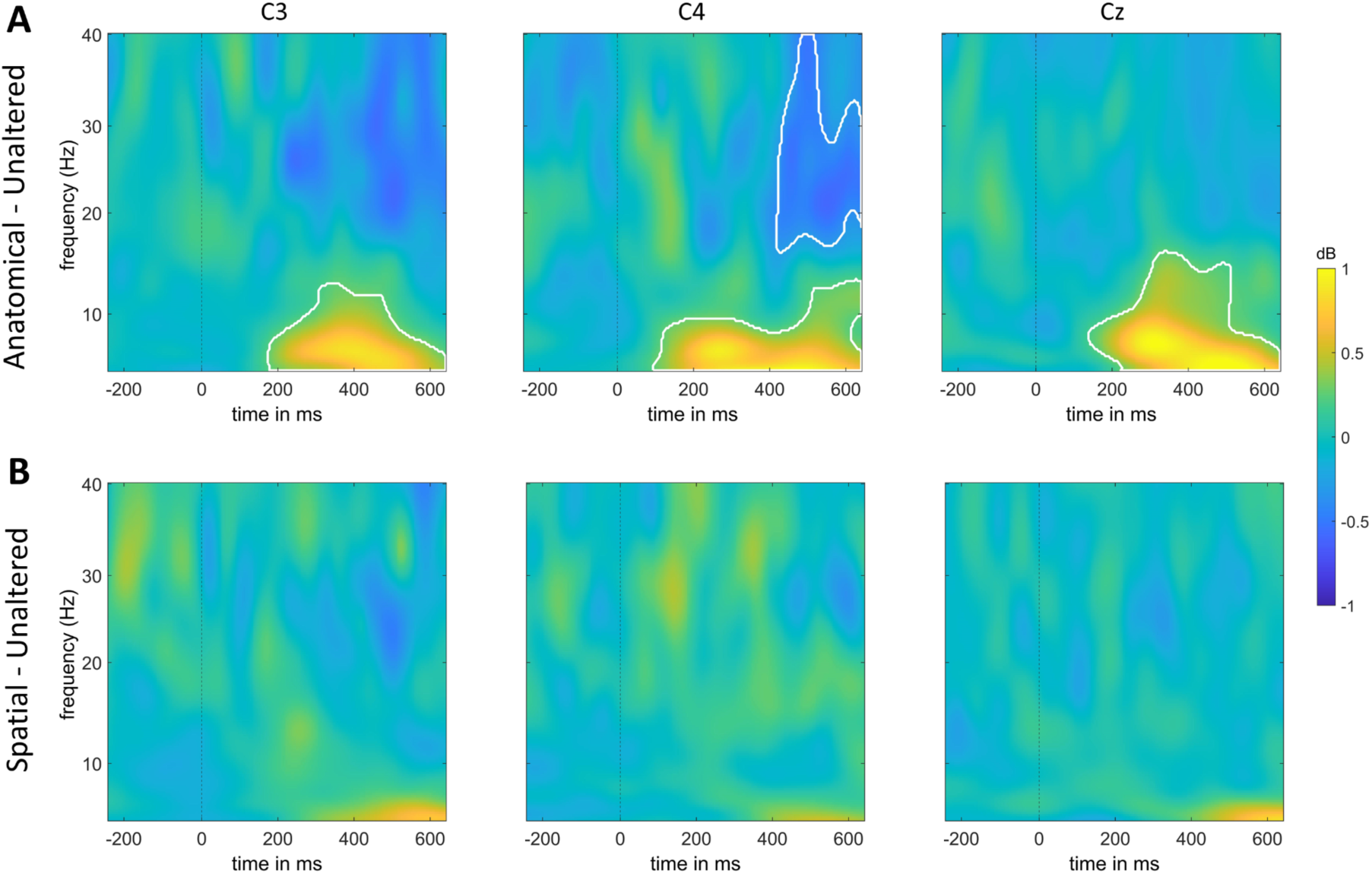
Average time-frequency power difference maps for pre-registered electrodes. **A**: Anatomical minus Unaltered. **B**: Spatial minus Unaltered. Significant clusters are marked with a white outline.

The results indicated that the difference in alpha frequency band activity was part of a larger activation difference patch, spanning lower frequencies as well (for the Anatomical condition, see Figure 3A). We then decided to explore EEG power modulations associated with SoA disruptions in the same ROI, for the same pre-registered electrodes (C3, C4 & Cz), in the theta frequency band (4-7 Hz). Results in this frequency band were very robust across all three electrodes, C3: t(29) = 4.8, p < 0.001, d = 0.89; C4: t(29) = 5.2, p < 0.001, d = 0.96; Cz: t(29) = 4.9, p < 0.001, d = 0.91 (see Figure 2). For the Spatial condition, we found a significant difference in theta frequency band in two (C3, C4) of our three electrodes of interest: C3: t(29) = 2.18, p < 0.05, d = 0.4; C4: t(29) = 1.97, p < 0.05, d = 0.37; Cz: t(29) = 1.49, p = 0.07, d = 0.27 (see Figure 2).

### Multivariate Pattern Analysis (MVPA)

To investigate the electrophysiological perturbations in response to SoA disruptions, we trained classifiers to distinguish between 1) Unaltered and Anatomical trials and 2) Unaltered and Spatial trials on both time-domain and frequency-transformed data. Above-chance classification implies that relevant information about the decoded stimulus feature is present in the neural data, meaning that some feature processing occurred (Hebart & Baker, 2018).

### Time-Domain

We first trained a classifier on time-domain data to distinguish between Unaltered trials and trials containing either one of the alterations (Anatomical or Spatial) separately. Consistent with the univariate approach, classification of the Anatomical condition revealed above-chance classification performance, with a classification accuracy ranging between 0.53 and 0.68 during the time period from 180ms to 1195ms after finger movement onset (average classification accuracy for our ROI was 0.65, average classification accuracy for the entire observed significant cluster was 0.61, p < 0.001, peak time: 460ms, see Figure 4A). All participants showed average above-chance area under the curve (AUC) values for the pre-registered time window (see supplementary Figure S2). For the Spatial condition, we found an above-chance classification performance with a classification accuracy ranging between 0.52 and 0.61 during the time period from 290ms to 1195ms after finger movement onset (average classification accuracy for our ROI was 0.56, average classification accuracy for the entire observed significant cluster was 0.57, p < 0.001, peak time: 752ms, see Figure 4A). All except three participants showed average above-chance AUC values for the pre-registered time window (see Supplementary Figure S2).

**Figure 4.**
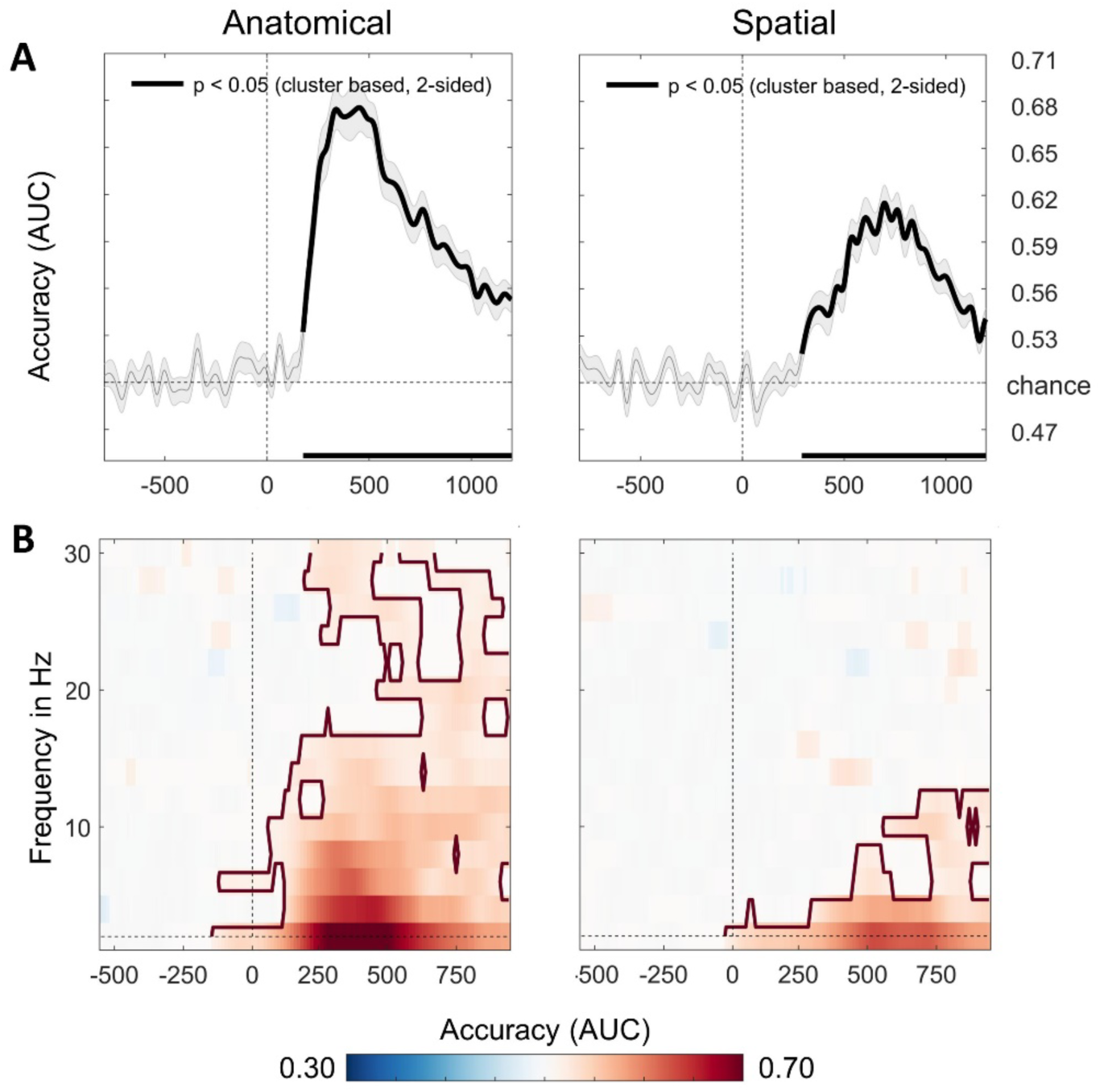
Multivariate classification of time-domain and frequency-transformed data. **A**: Multivariate classification accuracies of time-domain data resulting from comparing Unaltered vs. Anatomical (left) and Unaltered vs. Spatial (right). Bold lines within and at the bottom of the panels highlight significant differences (p < 0.05) under two-sided cluster- based permutation. Gray-shaded areas indicate the standard error of the mean across participants. **B**: Multivariate classification accuracies of frequency-transformed data resulting from comparing Unaltered vs. Anatomical (left) and Unaltered vs. Spatial (right). Classifier accuracies (AUC) are plotted across a time-frequency window of -500ms to 800ms and 2-30 Hz. Classifier accuracies are thresholded (cluster-based corrected, double-sided, p < 0.05), and significant clusters are outlined with a solid red line.

### Frequency-Domain

Classification of frequency-transformed data revealed a significant cluster in the Anatomical condition, widespread across time and frequency, spanning a time window of -140ms to 940ms, ranging from 2 to 30 Hz (p < 0.05, see Figure 4B). In line with the time-frequency analyses, the highest decoding accuracy was observed in the alpha and theta frequency bands (peak decoding: 370ms at 2 Hz). This finding is in line with our findings from the time-frequency analysis, showing the biggest contrast between Unaltered and Anatomical conditions around 350ms in low frequencies (see Figure 4B), as well as with previous literature (Bu-Omer et al., 2021; Kang et al., 2015; Savoie et al., 2018; Zama et al., 2019). In the Spatial condition, we found a significant above-chance classification cluster, ranging from 2 to 12 Hz, from -20ms to 940ms (p < 0.001, see Figure 4B). Similar to the Anatomical condition, the strongest decoding effect was observed in the lower frequencies (peak decoding: 520ms at 2 Hz). This again aligns with the time-frequency analysis, which showed the biggest contrast between Unaltered and Spatial conditions around 550ms in low frequencies (though significance was not reached; see Figure 3B). The finding that a multivariate approach captured a neural signature of the

Spatial SoA manipulation indicates that the effect might have been too subtle for the univariate time-frequency analysis to detect. It should also be noted that our classifier used data from all electrodes, in contrast to the time-frequency analysis, which only focused on the averages of pre-registered electrodes.

### Cross-decoding

Having established the presence of discriminatory neural activation patterns in both the Anatomical and Spatial contrasts, we then investigated the extent to which this information is shared between different conditions. The presence of shared information across conditions might indicate the existence of a domain-general SoA mechanism, rather than computations driven by specific sensorimotor conflicts. To address this question, we first trained a classifier on the Anatomical contrast (Unaltered vs. Anatomical) and tested it on the Spatial contrast (Unaltered vs. Spatial). Cross-classification of the time-domain data demonstrated a significant late cluster, with a classification accuracy ranging between 0.52 and 0.54 during the time period from 665ms to 730ms after movement onset (average classification accuracy = 0.53, p = 0.02, peak time: 688ms, see Figure 5A). Next, we performed the same cross-classification on frequency-transformed data to investigate the frequency characteristics of the shared information across conditions. This yielded an above-chance classification cluster, ranging from 2 to 6 Hz, from 440ms to 840ms (p < 0.001, peak time: 560ms at 2 Hz, see Figure 5B).

**Figure 5.**
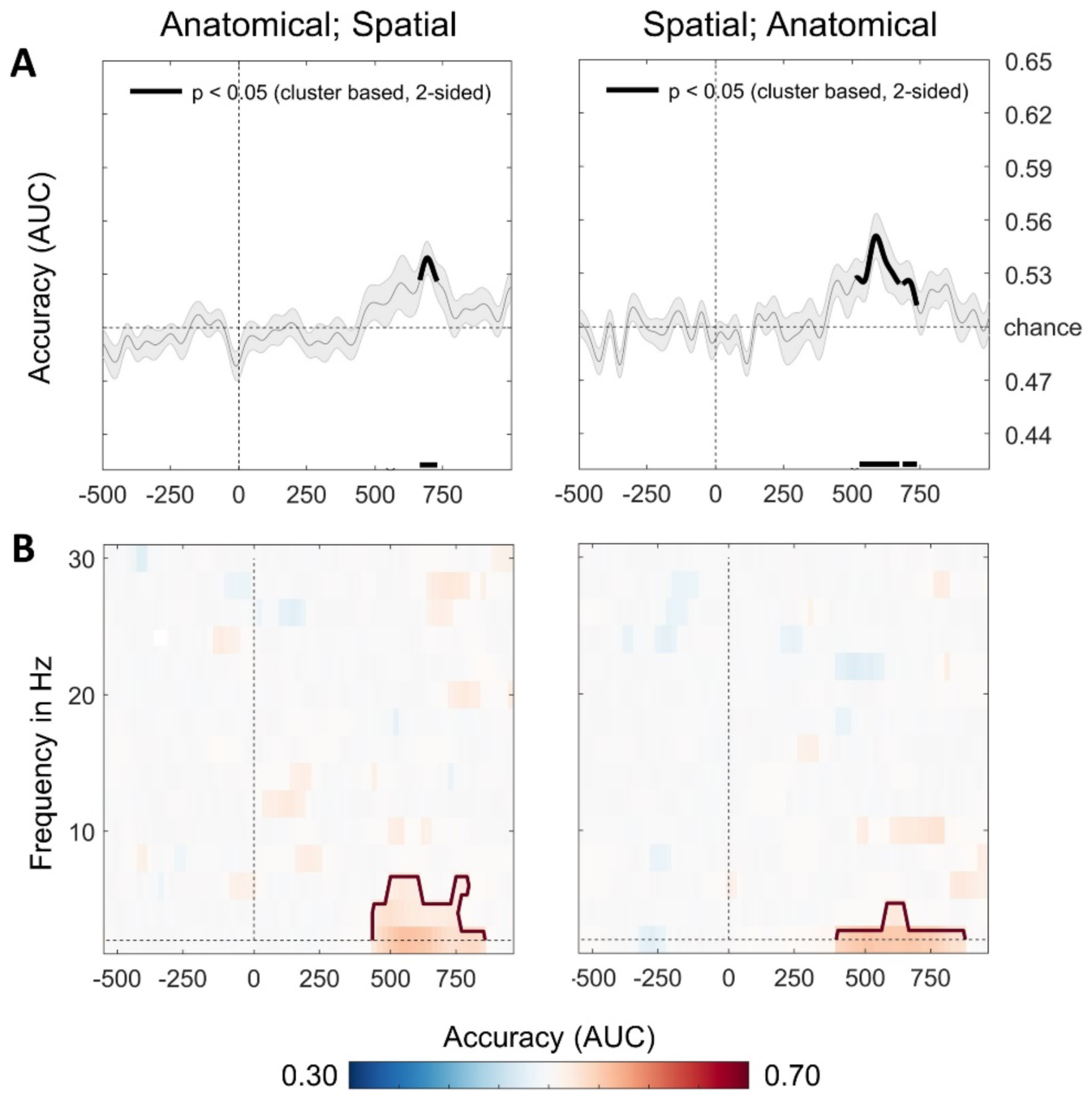
Cross-decoding multivariate classification accuracies. **A**: Multivariate classification accuracies of time-domain data resulting from training on Anatomical vs. Unaltered and testing on Spatial vs. Unaltered (left), and training on Spatial vs. Unaltered and testing on Anatomical vs. Unaltered (right). Bold lines within and at the bottom of the panels highlight significant differences (p < 0.05). Gray-shaded areas indicate the standard error of the mean across participants. **B**: Same as A, only for frequency-transformed data. Classifier accuracies of frequency data are plotted across a time-frequency window of -500ms to 800ms and 2-30 Hz. Classifier accuracies are thresholded (cluster-based corrected, double-sided, p < 0.05), significant clusters are outlined with a solid red line.

Subsequently, we trained a classifier on the Spatial contrast (Unaltered vs. Spatial) and tested it on the Anatomical contrast (Unaltered vs. Anatomical). Cross-classification of the time-domain data yielded two adjacent clusters. The first one had a classification accuracy ranging between 0.52 and 0.55 during the time period from 512ms to 672ms after movement onset (average classification accuracy = 0.54, p < 0.001, peak time: 580ms, see Figure 5A). The second one had a classification accuracy ranging between 0.53 and 0.54 during the time period from 684ms to 736ms after movement onset (average classification accuracy = 0.53, p < 0.021, peak time: 690ms, see Figure 5A). Next, we performed the same cross-classification on frequency-transformed data to investigate the frequency characteristics of the shared information across conditions. This yielded an above-chance classification cluster, ranging from 2 to 4 Hz, from 400ms to 860ms (p < 0.001, peak decoding: 500ms at 2 Hz, see Figure 5B).

## Discussion

In the current pre-registered study, we investigated the neural basis of embodied Sense of Agency. Utilizing a well-established paradigm involving a real-time virtual-reality hand model to manipulate agency, we demonstrated that reduced SoA is associated with increased alpha and theta power. Additionally, our findings provide electrophysiological evidence supporting the two-step model of agency.

Reduced SoA was consistently associated with decreased attenuation of activity in the alpha and theta frequency bands starting ∼180ms after movement onset in all three pre-registered electrodes in the Anatomical condition, as well as in C3 in the Spatial condition. Our findings are consistent with previous studies demonstrating that a larger power decrease in the alpha frequency band is associated with higher reported levels of agency (Bu-Omer et al., 2021; Kang et al., 2015), and are in line with the observed increase in alpha and low-beta power in response to sensory prediction errors over central electrodes (Savoie et al., 2018). While activity in the alpha frequency band is not a unitary phenomenon, alpha power synchronization, reflected by an increase in alpha power, is widely associated with functional inhibition (for reviews, see Jensen & Mazaheri, 2010; Klimesch et al., 2007). This inhibition indirectly facilitates the processing of incoming sensory information by suppressing irrelevant or distracting stimuli that might otherwise interfere with performance (Jensen & Mazaheri, 2010; Klimesch et al., 2007). Accordingly, an increase in alpha power in conditions characterized by a diminished SoA may be linked to the inhibition of conflicting information.

The observed increase in theta power with a reduced SoA can be explained in various ways. Parietal theta power has previously been associated with the processing of sensory prediction errors (Savoie et al., 2018) and has been found to scale with prediction error size (Arrighi et al., 2016). This is in line with the predictive coding theory, which proposes that feedforward prediction errors are carried by theta and gamma band synchronization and feedback communication by alpha and beta rhythms (Bastos et al., 2015; Halgren et al., 2019; Michalareas et al., 2016; Richter et al., 2018). Additionally, parietal theta band activity also seems related to motor adaptation in response to sensory prediction errors, where an increase in theta power has been observed in response to visual manipulations (Perfetti et al., 2011; Tzvi et al., 2022). These studies may explain why time-frequency results showed a strong finding for anatomical agency manipulations, yet none for spatial manipulations. The immediate, time-locked, and salient manipulation in the Anatomical condition may have elicited a larger prediction error, in contrast to the Spatial condition, where the manipulation unfolded over time and more evidence accumulation was required for a prediction error to occur. Additionally, the smaller prediction error might have led to a smaller motor adaptation response, which together could account for the lack of univariate time-frequency findings during Spatial agency violations.

Decoding of both types of agency manipulations demonstrated clear above-chance classification. Similar to the time pattern of the time-frequency analyses, reliable decoding started around 180ms after movement onset and was strongest in the alpha and theta frequency bands. It is particularly interesting that the Spatial condition demonstrated a large above-chance classification cluster, in contrast to the univariate time-frequency analysis, where the Spatial condition only yielded significant results in the ROI in a single electrode. A key difference between univariate and multivariate methods lies in the fact that the classifiers employed in decoding analyses utilize information that is averaged out in univariate analysis, making it a more sensitive method compared to traditional averaged-based techniques (Grootswagers et al., 2017). This could explain the difference in findings for the Spatial condition using MVPA and non-parametric permutation time-frequency analysis.

Our time-frequency and MVPA results are in line with the early sensorimotor computations of comparator mechanisms in the two-step agency model (Synofzik et al., 2008). To examine the involvement of higher-order mechanisms in the establishment of agency, as proposed by this model, we investigated whether neural patterns were shared between the two distinct types of agency manipulations. Shared neural patterns between the Anatomical and Spatial conditions would indicate that neural representations of reduced SoA are consistent across conditions, regardless of condition-specific sensorimotor conflicts, and hence, may point towards a domain-general SoA mechanism.

Our results demonstrate reliable cross-decoding from Unaltered vs. Anatomical to Unaltered vs. Spatial and vice versa, both in the time and frequency domain. These findings provide electrophysiological evidence for a late domain-general mechanism in SoA formation. As visual information differs across agency manipulations (see Figure 1B), the shared information across conditions is not likely to originate from similar low-level visual input. This conclusion is reinforced by the late emergence of the shared features, around 500ms in the time domain and 400ms in the frequency domain, pointing towards the involvement of a higher-order mechanism in the attribution of agency. Therefore, our findings support the idea that early SoA correlates are context-specific, while late higher-order mechanisms integrate these context-specific comparators.

The two-step model of agency (Synofzik et al., 2008) proposes early idiosyncratic information and late components of higher cognitive judgment. This model considers the information generated in the comparator mechanism (early, low-level step) to establish a conceptual Judgment of Agency (late, high-level step). In accordance with this account, we would expect early localized information specific to unique sensorimotor conflict manipulations, and a late component of domain-general SoA judgment. Our results show exactly that: Classification within each condition was significant, reaching peak classification accuracy early in the process (see Figure 4), while cross-classification was only significant later in the process (see Figure 5). We propose that the early component may reflect a computation of prediction error specific to the sensorimotor conflict, which explains why there is no shared information between the two contrasts early in the process. Later in the process, these specific prediction errors converge into a domain-general high-level SoA computation. To the best of our knowledge, we provide the first electrophysiological evidence to support the two-step account of SoA formation (Synofzik et al., 2008).

Our research has some limitations. First, the variability of participants’ confidence ratings was too small to perform any meaningful metacognition analysis. Second, to our surprise, a significant power difference when comparing Unaltered and Spatial trials was only found in a single electrode for alpha (C3, see Figure 2) and for none of the electrodes in high beta (See Supplementary Figure S1). This may be due to the higher sensitivity for anatomical prediction violations (see Daprati et al., 1997; Franck et al., 2001; Krugwasser et al., 2019). Indeed, the behavioral results indicate a substantially higher uncertainty in the Spatial condition compared to the Anatomical condition; notably, the variance is 23- fold higher (see Figure 1C). Lastly, the participants’ cohort was not equally divided in gender, as it contained twice as many females as males.

Looking forward, future research could explore the nuanced interactions between neural oscillations and SoA. One potential avenue for advancement lies in designing experiments that explore how different levels of sensorimotor alteration interact with various aspects of SoA. For example, investigating how subtle variations in sensory feedback influence SoA perception could provide deeper insights into the neural signature of agency. Another field to explore is the electrophysiological SoA-related activity in individuals with psychiatric conditions such as psychosis, who are known to have deficits in SoA (Harduf et al., 2023; Krugwasser et al., 2022; Raballo et al., 2011; Sass & Parnas, 2003). Examining the electrophysiological activity associated with SoA in these populations can uncover unique neural signatures and gain insights into the underlying mechanisms of agency perception in clinical contexts.

In conclusion, our findings provide compelling insights into the neural mechanisms associated with the experience of agency. Moreover, our research adds to the growing body of evidence highlighting the role of alpha and theta rhythms in processing prediction errors and updating perceptual predictions, which are tightly connected to SoA formation. Finally, using MVPA, the current research sheds light on the ongoing debate regarding whether the Sense of Agency arises from the early sensorimotor process or through a higher-order cognitive mechanism by showing early-stage differences and late-stage similarities between different conditions, providing novel evidence that supports the two-step formation of agency.

## Materials and Methods

### Participants

Forty-seven right-handed healthy participants were recruited through the university and within the local community through advertisements and received monetary compensation for their participation. All participants had normal or corrected-to-normal vision, had no diagnosed neurological or psychiatric conditions, and did not use psychiatric medication (based on self-report). All gave written informed consent to participate in the study, which was conducted under the Declaration of Helsinki and approved by the ethics committee of the Gonda Multidisciplinary Brain Research Center, Bar-Ilan University.

The aim of the study was to investigate the electrophysiological correlates of embodied SoA. We pre-registered exclusion criteria to ensure the quality of behavioral and neural data. Thus, participants who performed poorly during the experiment, did not finish the experiment, or had particularly noisy EEG recordings, were excluded from further analysis (see detailed criteria in the Methods section of the Supplementary materials). These criteria ensured that the participants did not deviate from the experimenter’s instructions and secured a sufficient number of trials for analysis. The final dataset consisted of 30 participants (average age = 24.7 ± 1.9; 20 females).

### Experimental setup and Procedure

The current study design is based on an experimental design used in previously published studies from our lab (Krugwasser et al., 2019, 2022; Stern et al., 2020, 2022), with small adjustments to allow EEG recordings. During the experiment, a virtual-reality model of the participant’s hand was displayed in real-time. This virtual hand allowed displaying visual feedback of movements that were manipulated with respect to the participant’s actual finger movements (see Figure 1A, B). A computer was used to conduct the experiment (Intel Core i7 processor and 16 GB of RAM), running custom-made software (SLabVR, built using Unity 5.6.1). It was displayed on a 24-inch Dell P2417H monitor at a 1920x1080 resolution. Motion tracking and virtual modeling of each participant’s right hand was visualized by a Leap Motion controller (Leap Motion Inc., San Francisco, CA), creating a virtual-reality model of the hand with real-time motion tracking (Krugwasser et al., 2019; Stern et al., 2020). Responses were given with a numeric keyboard. The intrinsic delay of the system is estimated at ∼80ms. The physical setup included a computer screen facing up, occluding participants’ real hand from view, with the Leap Motion controller positioned facing down underneath.

Participants were comfortably seated in a dimly lit room and placed their right hand, facing upward, directly beneath the Leap Motion controller, enabling them to see the virtual hand on the monitor, but not their real hand. They were instructed to fixate on the monitor during the entire experiment to reduce irrelevant ocular movement and blinking artifacts (see Figure 1A). Each trial started with the presentation of the virtual hand on the screen with a black fixation cross located at the tip of the index finger, signaling the participants not to move yet. After 600ms, the fixation cross turned red, indicating participants to make a single bending movement with their index finger, which resulted in visual feedback of the movement. The virtual hand disappeared after 1800ms, marking the end of the trial.

Visual feedback from the virtual hand movement was manipulated in two ways. In the Anatomical condition, the virtual hand conducted a comparable finger movement to the actual hand but with the middle finger instead of the index finger. In the Spatial condition, the virtual hand finger movement occurred with a 14-degree angular deviation from the actual finger movement (see Figure 1B). These manipulations were used in previous studies in our lab and resulted in a significant reduction in SoA for the vast majority of participants (Krugwasser et al., 2019, 2022; Stern et al., 2020).

To measure participants’ SoA and confidence, two questions were presented in Hebrew after each trial: “Was the movement I’ve seen congruent with the movement I’ve made?”, followed by “How confident are you regarding your previous answer?”. Using their left hand, participants were instructed to answer the first question by pressing the left key for no and the right key for yes. Participants were asked to rate their confidence on a continuous scale, ranging from “Not confident” (i.e., -3) to “Very confident” (i.e., 3) by pressing the left or right key in the desired direction and pressing enter (see Figure 1A).

The experiment consisted of 600 trials, divided into five blocks of 120 trials, preceded by an additional block of practice trials at the beginning of the experiment. Of these, 300 trials served as baseline trials, where the virtual hand made an identical movement to the participants’ actual hand (Unaltered condition). The other 300 trials were equally divided between manipulated conditions, resulting in 150 trials for each of the Anatomical and Spatial conditions, with a single alteration per trial (i.e., no combinations between alterations). The order of trials was randomized for each participant.

## Data analyses

Data are described as the average (±standard deviation).

### Statistical analysis of behavioral data

The first block of practice trials and all trials during which participants failed to move their finger or answer the response questions were excluded from all further analyses. Participants had an average of 292 (±8) valid trials in the Unaltered condition, 146 (±6) in the Anatomical condition, and 146 (±5) in the Spatial condition. SoA was calculated for each participant and condition by averaging responses to the agency question (where SoA=1 and no-SoA=0). To test for differences in SoA across different conditions, we used paired t-tests separately for each sensorimotor aspect. Sensitivity and bias were calculated separately for each aspect, then correlated and t-tested for differences (see Krugwasser et al., 2019). Confidence was calculated similarly, based on the response to the confidence question (without sensitivity and bias). Importantly, non-significant results were analyzed with equivalent Bayesian analysis to quantify evidence for the null effect.

### EEG acquisition and preprocessing

Scalp EEG was recorded with 32 channels using a wireless 64-electrode g.Nautilus research EEG headset, at a sampling rate of 250 Hz using the g.Recorder software (g.tec medical engineering GmbH, Austria). Electrodes were positioned according to the standard 10-20 system, with Cz positioned over the vertex of the participants’ heads. The continuous EEG data were imported into MATLAB (R2017b; The MathWorks, Inc., Natick, MA) and pre-processed offline using functions from the EEGLAB toolbox v2020.0 (Delorme & Makeig, 2004). EEG data were high-pass (0.5 Hz) and low-pass (40 Hz) filtered to remove slow drifts and high-frequency fluctuations (Cohen, 2014). Noisy channels were identified visually and interpolated (spherical interpolation), and manual visual artifact rejection was performed. The data were re-referenced to the average activity of all EEG scalp electrodes. Next, independent component analysis (ICA) was performed on the data, a technique that is commonly used to separate the data into maximally different components (Makeig et al., 1997). This method allows for the removal of artifactual components such as blinks, saccades, and muscular activity without having to delete entire epochs (Cohen, 2014). A component was removed based on one or more of the following criteria: Either its power spectrum did not show a 1/f distribution, or its topography demonstrated sources at the edges of the scalp, as these components often represent eye movement and muscle artifacts, or its time course presented spurious bursts of activity. Afterward, a second visual artifact rejection was performed to remove any remaining artifacts. Next, the data was epoched (-800ms to 1200ms), time-locked to finger movement onset, and baseline correction was applied using the pre-stimulus interval (- 400ms to -100ms). This baseline period was chosen to account for the temporal smearing from the wavelet convolution (Cohen, 2014). After preprocessing, an average of 245 (±28.6) trials were included in the analyses for the Unaltered condition, 124 (±14.3) trials for the Anatomical condition, and 124 (±13.7) trials for the Spatial condition.

### Time-frequency analysis

All time-frequency analyses were performed using functions implemented in the EEGLAB and Fieldtrip toolboxes (Oostenveld et al., 2010). Time-frequency transforms were computed for each electrode in the frequency range of 3 to 40 Hz (frequency resolution of 0.25 Hz) by employing complex Morlet wavelets (window size = 0.5 seconds) with the number of cycles increasing linearly with increasing frequency (from 3 cycles at the lowest to 8 cycles at the highest frequency), to account for the time-frequency trade-off (Cohen, 2014). The resulting event-related spectral perturbation (ERSP) measures are displayed in dB and represent a change in EEG power relative to the baseline. The epochs in each condition were averaged for each participant individually. To test the pre-registered hypothesis that the Anatomical and Spatial conditions are associated with a larger alpha attenuation, as well as a modulation in high beta power, we first performed analyses on predefined ROI, ranging from 100ms to 500ms, including motor electrodes C3, C4, and Cz for both the alpha and high beta frequency bands, subsequently.

These electrodes were chosen in light of previous research indicating their involvement in SoA-related processes (Kang et al., 2015; Shibuya et al., 2021). Next, average frequency power was extracted for each participant, ROI, and condition. To test for significance, we performed paired t-tests on the average frequency power per ROI (alpha and high beta) to compare the difference in power between the Unaltered condition and both the Anatomical and Spatial conditions. T-tests were one-tailed for analysis investigating the alpha frequency band and two-tailed for beta frequencies and explorative analyses. Alpha level was set to 5% across all analyses.

To statistically explore the comparison of the whole epoch power maps, we used a non-parametric cluster-based permutation approach implemented in FieldTrip (Maris & Oostenveld, 2007). In short, this well-established method first defines clusters by performing independent samples t-tests using subject-by-subject averages for each condition. After clusters in the original data were identified, 1000 random permutations were performed to create a permutation distribution of the largest clusters that were found after each permutation (cluster size was defined by summing t-values). Finally, the cluster statistic (the sum of the t-values) of the original cluster was computed as the proportion of random permutations that yielded a cluster statistic larger than the non-permuted cluster statistic (Maris & Oostenveld, 2007).

### Multivariate pattern analysis (MVPA)

In addition to the univariate approach, a multivariate backward decoding classification algorithm (linear discriminant analysis, LDA) was applied to the data using the open-source MATLAB-based Amsterdam Decoding and Modeling (ADAM) toolbox (version 1.05, see Fahrenfort et al., 2018). This was done because, unlike univariate average-based techniques, multivariate techniques take into account the relationships between multiple variables, resulting in higher sensitivity to subtle differences between conditions. Additionally, MVPA allows for the investigation of to what extent different agency violations share neural patterns (Grootswagers et al., 2017).

Classifiers were applied to both time-domain and frequency-transformed data (-800ms to 1200ms; 2-30 Hz). All 32 individual EEG electrodes were considered as features so that changes in topography, time, amplitude (for time-domain data), or power (for frequency-transformed data) were accounted for.

The classifier was trained and tested according to the within-subject classification approach separately on data from each participant, using a 10-fold cross-validation procedure to prevent overfitting (Wang et al., 2020). To this end, trials from each participant were divided into 10 equal-sized folds, as a classifier was trained on 9 folds and tested on the remaining fold. This procedure was repeated for every time point and every fold was used for testing once. Classification accuracy was averaged across folds.

To avoid classification bias in favor of the overrepresented condition, imbalances between the number of trials between conditions were balanced through oversampling (duplicating trials). Classification accuracy was assessed using AUC. Rather than averaging over binary class decisions, AUC takes into account the degree of confidence (as the distance from the decision boundary) when assigning a data point to a class, and is therefore considered a more sensitive, non-parametric, criterion-free measure of classification compared to ‘standard accuracy’ (Hand & Till, 2001). An AUC value of 0.5 indicates chance level performance, whereas a higher AUC value implies that relevant information is present in the data, allowing the distinction between classes.

Statistical significance was assessed across participants by testing AUC scores per time point against chance level performance of 0.5, with double-sided t-tests. Cluster-based permutation testing, as previously explained, was used to correct for multiple comparisons (p < 0.05, 1000 iterations). Significant above-chance level clusters imply that the neural patterns distinguishing between conditions are present. Note that this procedure should be interpreted as fixed effects (Allefeld et al., 2016), but it is nonetheless standard in the scientific community. Since spectral decoding can be time-consuming, the data were down-sampled to 100 Hz prior to spectral decoding and analyzed within the frequency range of 2 to 30 Hz, with increments of 2 Hz.

### Deviations From Pre-registration

Contrary to the analysis plan we pre-registered, we did not directly compare trials in which participants reported SoA to trials in which they did not feel SoA. The behavioral results indicated that participants had near perfect performance thus making this comparison redundant.

## Author Contribution

Amit Regev Krugwasser: Conceptualization, Methodology, Data collection, Data curation, Data pre-processing, Data Analysis, Visualization, Writing, Project administration. Reina van der Goot: Data pre-processing, Data Analysis, Visualization, Writing. Geffen Markusfeld: Data collection, Data curation, Data pre-processing. Yair Zvilichovsky: Software development. Roy Salomon: Conceptualization, Methodology, Data Analysis, Writing, Funding acquisition.

## References

Agnew, Z., & Wise, R. J. S. (2008). Separate Areas for Mirror Responses and Agency within the Parietal Operculum. The Journal of Neuroscience, 28(47), 12268–12273. 10.1523/JNEUROSCI.2836-08.2008

Allefeld, C., Görgen, K., & Haynes, J.-D. (2016). Valid population inference for information-based imaging: From the second-level t-test to prevalence inference. NeuroImage, 141, 378–392. 10.1016/j.neuroimage.2016.07.040

Andre, P., Cantore, N., Lucibello, L., Migliaccio, P., Rossi, B., Carboncini, M. C., Aloisi, A. M., Manzoni, D., & Arrighi, P. (2024). The cerebellum monitors errors and entrains executive networks. Brain Research, 1826, 148730. 10.1016/j.brainres.2023.148730

Applebaum, A., Netzer, O., Stern, Y., Zvilichovsky, Y., Mashiah, O., & Salomon, R. (2024). The Body Knows Better: Sensorimotor signals reveal “Suboptimal” inference of the Sense of Agency in the human mind (p. 2024.02.14.579431). bioRxiv. 10.1101/2024.02.14.579431

Arrighi, P., Bonfiglio, L., Minichilli, F., Cantore, N., Carboncini, M. C., Piccotti, E., Rossi, B., & Andre, P. (2016). EEG Theta Dynamics within Frontal and Parietal Cortices for Error Processing during Reaching Movements in a Prism Adaptation Study Altering Visuo- Motor Predictive Planning. PLOS ONE, 11(3), e0150265. 10.1371/journal.pone.0150265

Arzy, S., Thut, G., Mohr, C., Michel, C. M., & Blanke, O. (2006). Neural Basis of Embodiment: Distinct Contributions of Temporoparietal Junction and Extrastriate Body Area. The Journal of Neuroscience, 26(31), 8074–8081. 10.1523/JNEUROSCI.0745-06.2006

Assal, F., Schwartz, S., & Vuilleumier, P. (2007). Moving with or without will: Functional neural correlates of alien hand syndrome. Annals of Neurology, 62(3), 301–306.

Balslev, D., Nielsen, F. Å., Lund, T. E., Law, I., & Paulson, O. B. (2006). Similar brain networks for detecting visuo-motor and visuo-proprioceptive synchrony. NeuroImage, 31(1), 308–312. 10.1016/j.neuroimage.2005.11.037

Bastos, A. M., Vezoli, J., Bosman, C. A., Schoffelen, J.-M., Oostenveld, R., Dowdall, J. R., De Weerd, P., Kennedy, H., & Fries, P. (2015). Visual Areas Exert Feedforward and Feedback Influences through Distinct Frequency Channels. Neuron, 85(2), 390–401. 10.1016/j.neuron.2014.12.018

Blanke, O. (2012). Multisensory brain mechanisms of bodily self-consciousness. Nature Reviews Neuroscience, 13(8), Article 8. 10.1038/nrn3292

Braun, N., Debener, S., Spychala, N., Bongartz, E., Sörös, P., Müller, H. H. O., & Philipsen, A. (2018). The Senses of Agency and Ownership: A Review. Frontiers in Psychology, 9. 10.3389/fpsyg.2018.00535

Bu-Omer, H. M., Gofuku, A., Sato, K., & Miyakoshi, M. (2021). Parieto-Occipital Alpha and Low-Beta EEG Power Reflect Sense of Agency. Brain Sciences, 11(6), Article 6. 10.3390/brainsci11060743

Cohen, M. X. (2014). Analyzing Neural Time Series Data: Theory and Practice. MIT Press.

Daprati, E., Franck, N., Georgieff, N., Proust, J., Pacherie, E., Dalery, J., & Jeannerod, M. (1997). Looking for the agent: An investigation into consciousness of action and self-consciousness in schizophrenic patients. Cognition, 65(1), 71–86.

de Bézenac, C. E., Sluming, V., Gouws, A., & Corcoran, R. (2016). Neural response to modulating the probability that actions of self or other result in auditory tones: A parametric fMRI study into causal ambiguity. Biological Psychology, 119, 64–78. 10.1016/j.biopsycho.2016.07.003

Delorme, A., & Makeig, S. (2004). EEGLAB: An open source toolbox for analysis of single-trial EEG dynamics including independent component analysis. Journal of Neuroscience Methods, 134(1), 9–21. 10.1016/j.jneumeth.2003.10.009

Desantis, A., Roussel, C., & Waszak, F. (2014). The temporal dynamics of the perceptual consequences of action-effect prediction. Cognition, 132(3), 243–250. 10.1016/j.cognition.2014.04.010

Engbert, K., Wohlschläger, A., & Haggard, P. (2008). Who is causing what? The sense of agency is relational and efferent-triggered. Cognition, 107(2), 693–704.

Fahrenfort, J. J., van Driel, J., van Gaal, S., & Olivers, C. N. L. (2018). From ERPs to MVPA Using the Amsterdam Decoding and Modeling Toolbox (ADAM). Frontiers in Neuroscience, 12. 10.3389/fnins.2018.00368

Farrer, C., Frey, S. H., Van Horn, J. D., Tunik, E., Turk, D., Inati, S., & Grafton, S. T. (2008). The Angular Gyrus Computes Action Awareness Representations. Cerebral Cortex, 18(2), 254–261. 10.1093/cercor/bhm050

Franck, N., Farrer, C., Georgieff, N., Marie-Cardine, M., Daléry, J., d’Amato, T., & Jeannerod, M. (2001). Defective Recognition of One’s Own Actions in Patients With Schizophrenia. American Journal of Psychiatry, 158(3), 454–459. 10.1176/appi.ajp.158.3.454

Frewen, P., Schroeter, M. L., Riva, G., Cipresso, P., Fairfield, B., Padulo, C., Kemp, A. H., Palaniyappan, L., Owolabi, M., Kusi-Mensah, K., Polyakova, M., Fehertoi, N., D’Andrea, W., Lowe, L., & Northoff, G. (2020). Neuroimaging the consciousness of self: Review, and conceptual-methodological framework. Neuroscience & Biobehavioral Reviews, 112, 164–212. 10.1016/j.neubiorev.2020.01.023

Fukushima, H., Goto, Y., Maeda, T., Kato, M., & Umeda, S. (2013). Neural Substrates for Judgment of Self-Agency in Ambiguous Situations. PLOS ONE, 8(8), e72267. 10.1371/journal.pone.0072267

Gallagher, S. (2000). Philosophical conceptions of the self: Implications for cognitive science. Trends in Cognitive Sciences, 4(1), 14–21.

Gozli, D. G., & Brown, L. E. (2011). Agency and Control for the Integration of a Virtual Tool into the Peripersonal Space. Perception, 40(11), 1309–1319. 10.1068/p7027

Grootswagers, T., Wardle, S. G., & Carlson, T. A. (2017). Decoding dynamic brain patterns from evoked responses: A tutorial on multivariate pattern analysis applied to time series neuroimaging data. Journal of Cognitive Neuroscience, 29(4), 677–697. 10.1162/jocn_a_01068

Haggard, P. (2017). Sense of agency in the human brain. Nature Reviews Neuroscience, 18(4), 196–207. 10.1038/nrn.2017.14

Haggard, P., Clark, S., & Kalogeras, J. (2002). Voluntary action and conscious awareness. Nature Neuroscience, 5(4), 382–385. 10.1038/nn827

Halgren, M., Ulbert, I., Bastuji, H., Fabó, D., Erőss, L., Rey, M., Devinsky, O., Doyle, W. K., Mak-McCully, R., Halgren, E., Wittner, L., Chauvel, P., Heit, G., Eskandar, E., Mandell, A., & Cash, S. S. (2019). The generation and propagation of the human alpha rhythm. Proceedings of the National Academy of Sciences, 116(47), 23772–23782. 10.1073/pnas.1913092116

Han, N., Jack, B. N., Hughes, G., Elijah, R. B., & Whitford, T. J. (2021). Sensory attenuation in the absence of movement: Differentiating motor action from sense of agency. Cortex, 141, 436–448. 10.1016/j.cortex.2021.04.010

Hand, D. J., & Till, R. J. (2001). A Simple Generalisation of the Area Under the ROC Curve for Multiple Class Classification Problems. Machine Learning, 45(2), 171–186. 10.1023/A:1010920819831

Harduf, A., Panishev, G., Harel, E. V., Stern, Y., & Salomon, R. (2023). The bodily self from psychosis to psychedelics. Scientific Reports, 13(1), 21209. 10.1038/s41598-023-47600-z

Harduf, A., Shaked, A., Yaniv, A. U., & Salomon, R. (2023). Disentangling the Neural Correlates of Agency, Ownership and Multisensory Processing. NeuroImage, 120255. 10.1016/j.neuroimage.2023.120255

Hebart, M. N., & Baker, C. I. (2018). Deconstructing multivariate decoding for the study of brain function. NeuroImage, 180(Pt A), 4–18. 10.1016/j.neuroimage.2017.08.005

Horváth, J. (2015). Action-related auditory ERP attenuation: Paradigms and hypotheses. Brain Research, 1626, 54–65. 10.1016/j.brainres.2015.03.038

Hughes, G., & Waszak, F. (2011). ERP correlates of action effect prediction and visual sensory attenuation in voluntary action. NeuroImage, 56(3), 1632–1640. 10.1016/j.neuroimage.2011.02.057

Jensen, O., & Mazaheri, A. (2010). Shaping Functional Architecture by Oscillatory Alpha Activity: Gating by Inhibition. Frontiers in Human Neuroscience, 4. 10.3389/fnhum.2010.00186

Kang, S. Y., Im, C.-H., Shim, M., Nahab, F. B., Park, J., Kim, D.-W., Kakareka, J., Miletta, N., & Hallett, M. (2015). Brain Networks Responsible for Sense of Agency: An EEG Study. PLoS ONE, 10(8), e0135261. 10.1371/journal.pone.0135261

Karnath, H.-O., & Baier, B. (2010). Right insula for our sense of limb ownership and self-awareness of actions. Brain Structure and Function, 214(5–6), 411–417.

Klimesch, W., Sauseng, P., & Hanslmayr, S. (2007). EEG alpha oscillations: The inhibition-timing hypothesis. Brain Research Reviews, 53(1), 63–88. 10.1016/j.brainresrev.2006.06.003

Knoblich, G., Elsner, B., Aschersleben, G., & Metzinger, T. (2003). Grounding the self in action. Consciousness and Cognition, 12(4), 487–494.

Krugwasser, A. R., Harel, E. V., & Salomon, R. (2019). The boundaries of the self: The sense of agency across different sensorimotor aspects. Journal of Vision, 19(4), 14–14. 10.1167/19.4.14

Krugwasser, A. R., Stern, Y., Faivre, N., Harel, E. V., & Salomon, R. (2022). Impaired sense of agency and associated confidence in psychosis. Schizophrenia, 8(1), 32.

Kühn, S., Brass, M., & Haggard, P. (2013). Feeling in control: Neural correlates of experience of agency. Cortex, 49(7), 1935–1942. 10.1016/j.cortex.2012.09.002

Leptourgos, P., & Corlett, P. R. (2020). Embodied Predictions, Agency, and Psychosis. Frontiers in Big Data, 3, 27. 10.3389/fdata.2020.00027

Leube, D. T., Knoblich, G., Erb, M., Grodd, W., Bartels, M., & Kircher, T. T. J. (2003). The neural correlates of perceiving one’s own movements. NeuroImage, 20(4), 2084–2090. 10.1016/j.neuroimage.2003.07.033

Makeig, S., Jung, T.-P., Bell, A. J., Ghahremani, D., & Sejnowski, T. J. (1997). Blind separation of auditory event-related brain responses into independent components. Proceedings of the National Academy of Sciences, 94(20), 10979–10984. 10.1073/pnas.94.20.10979

Maris, E., & Oostenveld, R. (2007). Nonparametric statistical testing of EEG-and MEG-data. Journal of Neuroscience Methods, 164(1), 177–190.

Mas-Herrero, E., & Marco-Pallarés, J. (2014). Frontal Theta Oscillatory Activity Is a Common Mechanism for the Computation of Unexpected Outcomes and Learning Rate. Journal of Cognitive Neuroscience, 26(3), 447–458. 10.1162/jocn_a_00516

Michalareas, G., Vezoli, J., van Pelt, S., Schoffelen, J.-M., Kennedy, H., & Fries, P. (2016). Alpha-Beta and Gamma Rhythms Subserve Feedback and Feedforward Influences among Human Visual Cortical Areas. Neuron, 89(2), 384–397. 10.1016/j.neuron.2015.12.018

Miele, D. B., Wager, T. D., Mitchell, J. P., & Metcalfe, J. (2011). Dissociating Neural Correlates of Action Monitoring and Metacognition of Agency. Journal of Cognitive Neuroscience, 23(11), 3620–3636. 10.1162/jocn_a_00052

Mifsud, N. G., Beesley, T., Watson, T. L., & Whitford, T. J. (2016). Attenuation of auditory evoked potentials for hand and eye-initiated sounds. Biological Psychology, 120, 61–68. 10.1016/j.biopsycho.2016.08.011

Moore, J. W. (2016). What Is the Sense of Agency and Why Does it Matter? Frontiers in Psychology, 7. 10.3389/fpsyg.2016.01272

Moore, J. W., & Fletcher, P. C. (2012). Sense of agency in health and disease: A review of cue integration approaches. Consciousness and Cognition, 21(1), 59–68.

Moro, V., Pernigo, S., Scandola, M., Mainente, M., Avesani, R., & Aglioti, S. M. (2015). Contextual bottom-up and implicit top-down modulation of anarchic hand syndrome: A single-case report and a review of the literature. Neuropsychologia, 78, 122–129.

Nakashima, R. (2019). Beyond one’s body parts: Remote object movement with sense of agency involuntarily biases spatial attention. Psychonomic Bulletin & Review, 26(2), 576–582. 10.3758/s13423-018-1552-4

Newson, J. J., & Thiagarajan, T. C. (2019). EEG Frequency Bands in Psychiatric Disorders: A Review of Resting State Studies. Frontiers in Human Neuroscience, 12. https://www.frontiersin.org/articles/10.3389/fnhum.2018.00521

Ohata, R., Asai, T., Kadota, H., Shigemasu, H., Ogawa, K., & Imamizu, H. (2020). Sense of Agency Beyond Sensorimotor Process: Decoding Self-Other Action Attribution in the Human Brain. Cerebral Cortex, 30(7), 4076–4091. 10.1093/cercor/bhaa028

Oostenveld, R., Fries, P., Maris, E., & Schoffelen, J.-M. (2010). FieldTrip: Open source software for advanced analysis of MEG, EEG, and invasive electrophysiological data. Computational Intelligence and Neuroscience, 2011.

Park, H.-D., Bernasconi, F., Salomon, R., Tallon-Baudry, C., Spinelli, L., Seeck, M., Schaller, K., & Blanke, O. (2018). Neural sources and underlying mechanisms of neural responses to heartbeats, and their role in bodily self-consciousness: An intracranial EEG study. Cerebral Cortex, 28(7), 2351–2364.

Perfetti, B., Moisello, C., Landsness, E. C., Kvint, S., Lanzafame, S., Onofrj, M., Rocco, A. D., Tononi, G., & Ghilardi, M. F. (2011). Modulation of Gamma and Theta Spectral Amplitude and Phase Synchronization Is Associated with the Development of Visuo- Motor Learning. Journal of Neuroscience, 31(41), 14810–14819. 10.1523/JNEUROSCI.1319-11.2011

Pezzetta, R., Nicolardi, V., Tidoni, E., & Aglioti, S. M. (2018). Error, rather than its probability, elicits specific electrocortical signatures: A combined EEG-immersive virtual reality study of action observation. Journal of Neurophysiology, 120(3), 1107–1118. 10.1152/jn.00130.2018

Pyasik, M., Furlanetto, T., & Pia, L. (2019). The Role of Body-Related Afferent Signals in Human Sense of Agency. Journal of Experimental Neuroscience, 13, 1179069519849907. 10.1177/1179069519849907

Quesque, F., & Brass, M. (2019). The Role of the Temporoparietal Junction in Self-Other Distinction. Brain Topography, 32(6), 943–955. 10.1007/s10548-019-00737-5

Raballo, A., Sæbye, D., & Parnas, J. (2011). Looking at the Schizophrenia Spectrum Through the Prism of Self-disorders: An Empirical Study. Schizophrenia Bulletin, 37(2), 344–351. 10.1093/schbul/sbp056

Renes, R. A., van Haren, N. E. M., Aarts, H., & Vink, M. (2015). An exploratory fMRI study into inferences of self-agency. Social Cognitive and Affective Neuroscience, 10(5), 708–712. 10.1093/scan/nsu106

Richter, C. G., Coppola, R., & Bressler, S. L. (2018). Top-down beta oscillatory signaling conveys behavioral context in early visual cortex. Scientific Reports, 8(1), 6991. 10.1038/s41598-018-25267-1

Rondi-Reig, L., Paradis, A.-L., Lefort, J. M., Babayan, B. M., & Tobin, C. (2014). How the cerebellum may monitor sensory information for spatial representation. Frontiers in Systems Neuroscience, 8. 10.3389/fnsys.2014.00205

Sass, L. A., & Parnas, J. (2003). Schizophrenia, Consciousness, and the Self. Schizophrenia Bulletin, 29(3), 427–444. 10.1093/oxfordjournals.schbul.a007017

Savoie, F.-A., Thénault, F., Whittingstall, K., & Bernier, P.-M. (2018). Visuomotor Prediction Errors Modulate EEG Activity Over Parietal Cortex. Scientific Reports, 8(1), 12513. 10.1038/s41598-018-30609-0

Seghezzi, S., & Zapparoli, L. (2020). Predicting the Sensory Consequences of Self-Generated Actions: Pre-Supplementary Motor Area as Supra-Modal Hub in the Sense of Agency Experience. Brain Sciences, 10(11), Article 11. 10.3390/brainsci10110825

Seghezzi, S., Zirone, E., Paulesu, E., & Zapparoli, L. (2019). The Brain in (Willed) Action: A Meta-Analytical Comparison of Imaging Studies on Motor Intentionality and Sense of Agency. Frontiers in Psychology, 10. https://www.frontiersin.org/articles/10.3389/fpsyg.2019.00804

Serino, A., Bockbrader, M., Bertoni, T., Colachis IV, S., Solcà, M., Dunlap, C., Eipel, K., Ganzer, P., Annetta, N., Sharma, G., Orepic, P., Friedenberg, D., Sederberg, P., Faivre, N., Rezai, A., & Blanke, O. (2022). Sense of agency for intracortical brain–machine interfaces. Nature Human Behaviour, 6(4), Article 4. 10.1038/s41562-021-01233-2

Shibuya, S., Unenaka, S., Shimada, S., & Ohki, Y. (2021). Distinct modulation of mu and beta rhythm desynchronization during observation of embodied fake hand rotation. Neuropsychologia, 159, 107952. 10.1016/j.neuropsychologia.2021.107952

Spengler, S., von Cramon, D. Y., & Brass, M. (2009). Was it me or was it you? How the sense of agency originates from ideomotor learning revealed by fMRI. NeuroImage, 46(1), 290–298. 10.1016/j.neuroimage.2009.01.047

Stern, Y., Ben-Yehuda, I., Koren, D., Zaidel, A., & Salomon, R. (2022). The dynamic boundaries of the Self: Serial dependence in the Sense of Agency. Cortex, 152, 109–121.

Stern, Y., Koren, D., Moebus, R., Panishev, G., & Salomon, R. (2020). Assessing the relationship between sense of agency, the bodily-self and stress: Four virtual-reality experiments in healthy individuals. Journal of Clinical Medicine, 9(9), 2931.

Synofzik, M., Thier, P., Leube, D. T., Schlotterbeck, P., & Lindner, A. (2010). Misattributions of agency in schizophrenia are based on imprecise predictions about the sensory consequences of one’s actions. Brain, 133(1), 262–271.

Synofzik, M., Vosgerau, G., & Newen, A. (2008). Beyond the comparator model: A multifactorial two-step account of agency. Consciousness and Cognition, 17(1), 219–239.

Synofzik, M., Vosgerau, G., & Voss, M. (2013). The experience of agency: An interplay between prediction and postdiction. Frontiers in Psychology, 4, 127. Pmc. 10.3389/fpsyg.2013.00127

Tapal, A., Oren, E., Dar, R., & Eitam, B. (2017). The Sense of Agency Scale: A Measure of Consciously Perceived Control over One’s Mind, Body, and the Immediate Environment. Frontiers in Psychology, 8. 10.3389/fpsyg.2017.01552

Tsakiris, M., Haggard, P., Franck, N., Mainy, N., & Sirigu, A. (2005). A specific role for efferent information in self-recognition. Cognition, 96(3), 215–231.

Tsakiris, M., Longo, M. R., & Haggard, P. (2010). Having a body versus moving your body: Neural signatures of agency and body-ownership. Neuropsychologia, 48(9), 2740–2749. 10.1016/j.neuropsychologia.2010.05.021

Tzur, G., & Berger, A. (2007). When things look wrong: Theta activity in rule violation. Neuropsychologia, 45(13), 3122–3126. 10.1016/j.neuropsychologia.2007.05.004

Tzvi, E., Gajiyeva, L., Bindel, L., Hartwigsen, G., & Classen, J. (2022). Coherent theta oscillations in the cerebellum and supplementary motor area mediate visuomotor adaptation. NeuroImage, 251, 1–11. 10.1016/j.neuroimage.2022.118985

Villa, R., Ponsi, G., Scattolin, M., Panasiti, M. S., & Aglioti, S. M. (2022). Social, affective, and non-motoric bodily cues to the Sense of Agency: A systematic review of the experience of control. Neuroscience & Biobehavioral Reviews, 142, 104900. 10.1016/j.neubiorev.2022.104900

Villa, R., Tidoni, E., Porciello, G., & Aglioti, S. M. (2018). Violation of expectations about movement and goal achievement leads to Sense of Agency reduction. Experimental Brain Research, 236(7), 2123–2135. 10.1007/s00221-018-5286-3

Wang, Q., Cagna, B., Chaminade, T., & Takerkart, S. (2020). Inter-subject pattern analysis: A straightforward and powerful scheme for group-level MVPA. NeuroImage, 204. 10.1016/j.neuroimage.2019.116205

Wegner, D. M., & Wheatley, T. (1999). Apparent mental causation: Sources of the experience of will. American Psychologist, 54(7), 480.

Wen, W., Charles, L., & Haggard, P. (2023). Metacognition and sense of agency. Cognition, 241, 105622.

Wolpert, D. M., Ghahramani, Z., & Jordan, M. I. (1995). An internal model for sensorimotor integration. Science, 269(5232), 1880.

Yomogida, Y., Sugiura, M., Sassa, Y., Wakusawa, K., Sekiguchi, A., Fukushima, A., Takeuchi, H., Horie, K., Sato, S., & Kawashima, R. (2010). The neural basis of agency: An fMRI study. NeuroImage, 50(1), 198–207. 10.1016/j.neuroimage.2009.12.054

Zaidel, A., & Salomon, R. (2023). Multisensory decisions from self to world. Philosophical Transactions of the Royal Society B: Biological Sciences, 378(1886), 20220335. 10.1098/rstb.2022.0335

Zama, T., Takahashi, Y., & Shimada, S. (2019). Simultaneous EEG-NIRS Measurement of the Inferior Parietal Lobule During a Reaching Task With Delayed Visual Feedback. Frontiers in Human Neuroscience, 13. https://www.frontiersin.org/article/10.3389/fnhum.2019.00301

Zito, G. A., Wiest, R., & Aybek, S. (2020). Neural correlates of sense of agency in motor control: A neuroimaging meta-analysis. PLOS ONE, 15(6), e0234321. 10.1371/journal.pone.0234321

