## Supplementary Materials for "Neural Correlates of the Embodied Sense of Agency"

### Methods

***Participants.*** The following pre-registered exclusion criteria were used to ensure the quality of behavioral and neural data:

1. Participants with less than 75% self-attribution in Unaltered trials.
2. Participants who showed less than a 30% decrease of SoA between the no alteration condition and the altered conditions, calculated separately between conditions (Anatomical and Spatial).
3. Participants who were unable to comply with the task or failed to complete at least 80% of the experiment.
4. Participants that needed interpolation on 10 or more EEG channels.
5. After EEG pre-processing, participants whose number of trials fell below 60% of the total number of trials in the experiment (i.e. less than 180 trials in the Unaltered condition and 90 trials in the Anatomical/Spatial condition).

***Experimental set-up***. The background color of all the screens in this experiment was gray - RGB(127, 127, 127) - the text color was black (except the message indicating breaks, which was white), and the text font was Segoe UI.

***EEG acquisition.*** The locations of the EEG electrodes were: 'FP1', 'FP2', 'FZ', 'F3', 'F4', 'F7', 'F8', 'AF3', 'AF4', 'FC1', 'FC2', 'FC5', 'FC6', 'C3', 'CZ', 'C4', 'CP1', 'CP2', 'CP5', 'CP6', 'PZ', 'P3', 'P4', 'P7', 'P8', 'PO3', 'PO4', 'PO7',' PO8', 'OZ', 'T7' and 'T8'.

### Results

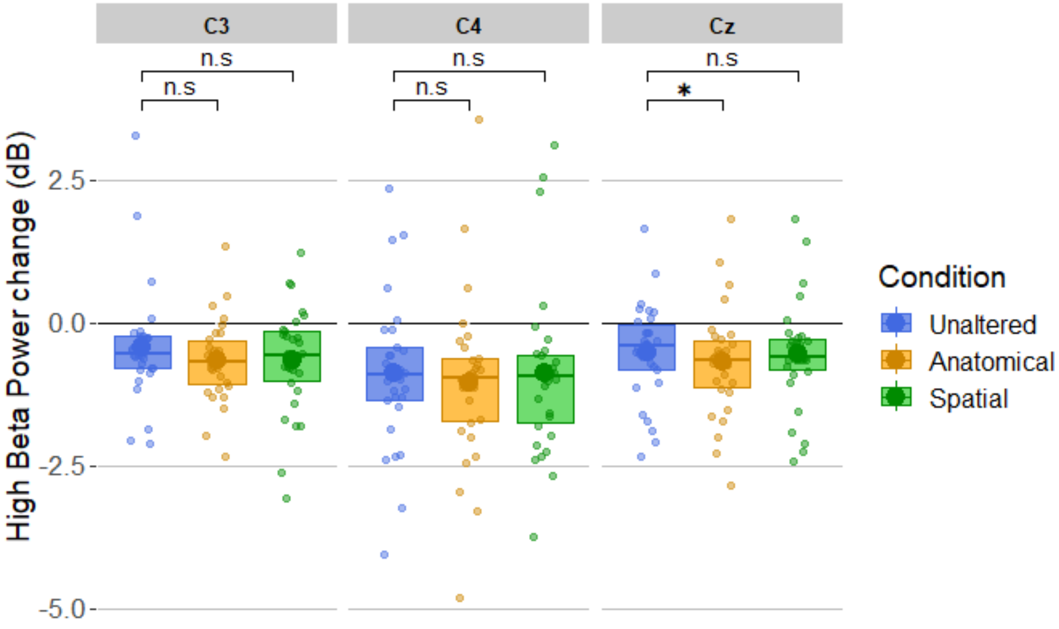

**Supplementary Figure S1.** Average (large circles) and individual (dots) power modulation for each electrode (C3, C4, Cz) in each condition (Unaltered, Anatomical, Spatial), in high beta frequency band, compared to baseline.

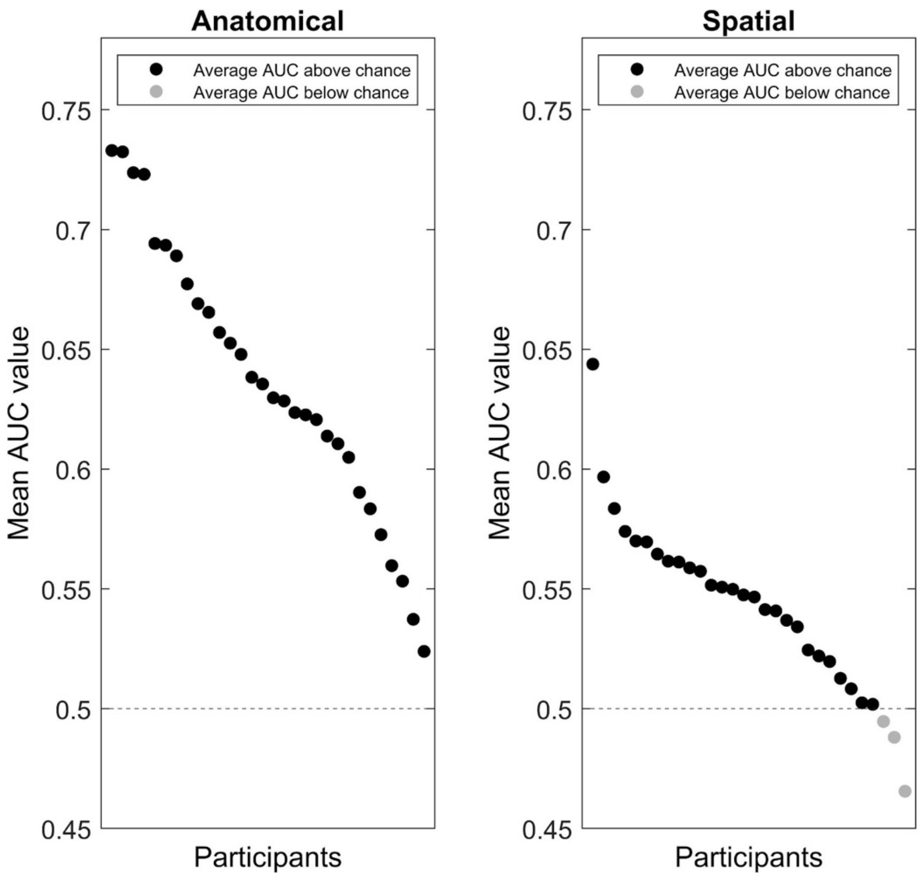


**Supplementary Figure S2.** Average AUC values for the pre-registered decoding time window (150-600 ms after finger movement onset) per participant for the Anatomical (left) and Spatial (right) conditions.

| | Unaltered | Anatomical | Spatial |
|---|---|---|---|
| Subjective SoA rating | 0.94 (0.06) | 0.013 (0.015) | 0.091 (0.1) |
| Confidence | 2.41 (0.5) | 2.76 (0.4) | 2.36 (0.55) |

**Supplementary Table S1.** Average (and SD) ratings of SoA and Confidence.
